# Parallel mechanisms of visual memory formation across distinct regions of the honey bee brain

**DOI:** 10.1101/2020.02.10.939355

**Authors:** Arián Avalos, Ian M. Traniello, Eddie Perez Claudio, Tugrul Giray

## Abstract

Visual learning is vital to the behavioral ecology of the Western honey bee (*Apis mellifera*). Honey bee workers forage for floral resources, a behavior that requires the learning and long-term memory of visual landmarks, but how these memories are mapped to the brain remains poorly understood. To address this gap in our understanding, we collected bees that successfully learned visual associations in a conditioned aversion paradigm and compared gene expression correlates of memory formation in the mushroom bodies, a higher-order sensory integration center classically thought to contribute to learning, as well as the optic lobes, the primary visual neuropil responsible for sensory transduction of visual information. We quantitated expression of *CREB* and *CaMKii*, two classical genetic markers of learning and *fen-1*, a gene specifically associated with punishment learning in vertebrates. As expected, we report substantial involvement of the mushroom bodies for all three markers but additionally demonstrate an unexpected involvement of the optic lobes across a similar time course. Our findings imply the molecular involvement of a sensory neuropil during visual associative learning parallel to a higher-order brain region, furthering our understanding of how a tiny brain processes environmental signals.

## Introduction

Visual learning is necessary for the survival and growth of honey bee societies. Honey bee foragers, bees that locate and gather resources for the colony, use visual cues to perform orientation flights (Cartwright and Collett, 1983; Menzel et al., 2005), locate floral resources (Chittka and Raine, 2006; Giurfa et al., 1995; Lehrer et al., 1995; Sen Sarma et al., 2010), and possibly to assess and select new nest sites (Seeley and Visscher, 2003; Visscher, 2007). Despite its importance, the neuroanatomical basis of visual memory formation remains poorly understood. We set out to address this issue by exploring the spatial and temporal aspects of learning in anatomically distinct sensory- and learning-associated regions of the honey bee brain.

The honey bee brain consists of ∼1,000,000 neurons, about 340,000 of which are Kenyon cells composing mushroom bodies (MB), centers of sensory integration, learning and memory (Heisenberg, 1998; Strausfeld, 2002; Witthöft, 1967). Visual stimulus perception and signal transduction are carried by adjacent, distinctly compartmentalized regions, the optic lobes (OL), which project visual input to the MB. By capitalizing on the contrast of cellular function in these discrete anatomic subunits, we can begin to understand the path of signal transduction of visual environmental information to sensory integration and processing and test the hypothesis that sensory regions may themselves store memories. To this end, we utilized expression analysis of established learning and memory genes in the primary visual neuropil and a higher-order processing center following a visual learning event.

In honey bees, long term memory (LTM) formation has been best characterized using classical conditioning of the proboscis extension response (PER) to olfactory stimuli (Bitterman et al., 1983; Menzel, 1999), with similar memory phases described following aversive conditioning (Agarwal et al., 2011; Nouvian and Galizia, 2019). Like in other behavioral systems, LTM in honey bees involves activation of the calcium/calmodulin kinases (CaMK) (Kamikouchi et al., 2000; Perisse et al., 2009). Phosphorylation of CaMKII leads to the subsequent activation of cyclic AMP response element-binding protein (CREB), which becomes active and modulates transcription for the long-term maintenance of newly formed associations (Eisenhardt et al., 2003; Kamikouchi et al., 2000; Kandel, 2001; Kandel, 2012; Matsumoto and Mizunami, 2002). The downstream activation of CREB via phosphorylation induces its function as a transcription factor in LTM processes (Bito et al., 1996; Kandel, 2001; Kandel, 2012; Lakhina et al., 2015). Our approach used expression profiles of *CaMKII* and *CREB*, established markers of the LTM process in the honey bee and other systems (Eisenhardt et al., 2003; Kamikouchi et al., 2000; Kandel, 2001; Kandel, 2012; Matsumoto et al., 2009; Menzel and Giurfa, 2001; Pasch et al., 2011; Perisse et al., 2009). Therefore, detectable changes in the expression of these genes following learning can serve as a predictive indicator for downstream LTM formation. In addition to established targets of the LTM process (*CaMKII, CREB*) we also explored the expression profile of the gene *flap structure-specific endonuclease 1* (*fen-1*). Though not yet described in honey bees, *fen-1* has been previously associated with aversive conditioning in vertebrate models (Saavedra-Rodríguez et al., 2009; Wang et al., 2003), thus making it a promising target gene for comparative analyses of aversive conditioning in the honey bee brain.

Considering both target expression and neural pathway input, we hypothesize that: 1) gene expression differences will be present in the MB alone, as mechanisms underlying memory formation may not be necessary in regions of sensory signal transduction; or, alternatively, 2) gene expression will follow the anatomical pathway visual input must take from sensory signal transduction to higher-order sensory processing with OL signal detected earlier than MB. By analyzing brain region-specific gene expression following a visual retention task, we begin to inform an understanding of the spatial and temporal dynamics of learning and memory.

## Materials and Methods

### Collections

We collected returning foraging worker honey bees at the research apiary at Gurabo Agricultural Research Station of the University of Puerto Rico in Gurabo, Puerto Rico. All workers were collected during peak foraging hours (8:00-17:00) (Mattu et al., 2012) by blocking the colony entrance with a wire mesh screen (6.32 mm^2^ aperture), then using a modified collection vacuum (Model 5911, Type 1, 12V DC; BioQuip, Rancho Dominguez, CA) to safely aspirate workers into a collection vessel. Immediately following collection, the wire mesh was removed, and the collection cage was extracted from the vacuum and sealed. Collected bees were provided with 50% sugar solution and transported to our research laboratory at the University of Puerto Rico, San Juan. Foragers were quickly placed in a rearing cage (Bug Dorm 1 Rearing Cage; BioQuip, Rancho Dominguez, CA) with food provided *ad libitum* and left overnight in a dark incubator set at 34°C.

### Electric Shock Avoidance assay

We used the Electric Shock Avoidance (ESA) assay to examine color-learning. This assay is a free-operant experimental paradigm that selectively isolates visual learning (Agarwal et al., 2011; Avalos et al., 2017). Foraging workers collected and transported to our research laboratory were quickly placed in a rearing cage one day following collection, and groups of 10 bees were sequentially extracted for the ESA (Avalos et al., 2017).

A simplified version of the ESA assay was used to train individual bees. The protocol presented the color using a Styrofoam™ block with equal halves of its surfaced lined with blue and yellow construction paper. The cassette was placed on top of the block during shock presentation, aligning the electrified area with the selected color of the grid. During recovery periods between the 5 min trials and short-term memory test, the cassette was placed in a dark incubator. Both color and position of shock were counterbalanced between groups of bees trained to avoid spatial learning independent of color.

Four experimental groups were sampled in this study: Naïve control (N_C_), Context control (C_C_), Shock control (S_C_), and Learned (L_E_). N_C_ bees were collected directly from the colony, acclimated in the incubator and anesthetized with a 15 s exposure to CO_2_ and flash-frozen 15 min later, upon recovery. This group therefore controls for baseline gene expression of a honey bee forager during experimental handling.

L_E_ bees were exposed to the same handling process as N_C_ bees, but, following recovery from CO_2_, were then subjected to training. In the training assay, we paired one of two colors with a mild shock (CS^+^) over two 5 min trial presentations. These two presentations were separated by a 10 min inter-trial interval (ITI) spent in a dark incubator to remove visual stimuli and avoid possible memory extinction in the absence of shock. Following training, L_E_ bees were again placed in the incubator for 20 min, then exposed to a one min short term memory (STM) test in which color but no shock was provided. C_C_ group bees went through the same process as learning group bees, but during the 5 min trials no shock was provided to either side. This group therefore provides a control of potential effects from bees being placed in the training arena.

For the S_C_ group, we also assayed 10 individuals at a time. We used a yoked control design in which one bee was designated “master” and experienced the same training as the L_E_ individuals, while the remaining nine bees were designated “yoked,” experiencing the same proportion of shock events and duration as the master bee but disassociated from the visual stimulus. In this way S_C_ individuals served as controls experiencing noxious stimuli in absence of color context.

For all groups, behavioral response was measured and cataloged by two observers via scan sampling. One observer scanned the grid every 15 seconds and conveyed the presence/absence of each bee on the shock side of the apparatus to the cataloguing participant. Response data were used to categorize individuals (*see below*). Individual bees were collected and flash frozen in liquid nitrogen immediately following recovery (N_C_ group) or at 20 or 80 min following the last presentation trial (all other groups). Individuals were kept at-80°C to await sample selection and gene expression analysis.

### Sample selection for molecular analysis

Across all samples, we screened for survival of handling (all), adequate interaction with the arena (C_C_, S_C_), and in the case of the L_E_ group, association of shock stimulus with color. To be suitable for gene expression analysis, all individuals needed to have survived handling and CO_2_ anesthetization. For those groups experiencing the apparatus, they were required to have shuttled between color regions at least three times. For L_E_ we additionally required that 1) they spend more than 50% of the last 2.5 min of the second trial on the safe side, and 2) on two of the possible four 2.5 min time blocks (*described below*). This selection scheme allowed us to identify bees that correctly associated shock with color and therefore experience learning. Criterion 1 focused on improvement: better-than-average performance in this time period suggests retention of learned information (Agarwal et al., 2011; Avalos et al., 2017). Criterion 2 assured that acquisition occurred throughout the assay. These additional criteria in L_E_ assured we identified bees that correctly formed an association between color and punishment, i.e. learned avoidance (Fig. S1). Any bee not meeting selection criteria was excluded from further analysis resulting in the following per-group sample sizes: 6 (N_C_), 16 (C_C_), 24 (S_C_), 18 (L_E_).

### Gene expression analysis

Head capsules were chipped on dry ice to expose the brain, glands, and optic lobe pigment, and the whole head was submerged in RNAlater^®^ ICE (Thermo Fisher Scientific, Waltham, MA) at −20°C for 16 hr (Fig. S2A). Brains were fully extracted on wet ice and regions of interest were dissected (Fig. S2B). We performed region-specific analysis aided by the honey bee brain atlas (Brandt et al., 2005; Rybak et al., 2010), dissecting out the MB and OL specifically. We also utilized the remaining tissue, composed of the protocerebrum, subesophageal ganglion (sEG), and AL as a conglomerate we reference here as central brain (CB) (Fig. S2C). We used the CB as a contrasting physiological control as they contain regions likely involved with signal transduction during visual LTM (e.g. protocerebrum) but also those not (e.g. sEG, AL). Regions were re-suspended in RNAlater^®^ ICE solution for later analysis. To obtain sufficient genetic material for analysis, we pooled brain regions from two individual bees randomly chosen from each behavioral group. This resulted in the final per-individual sample sizes of: N_C_ n = 3, C_C_ 20 min n = 3, C_C_ 80 min n = 5, S_C_ 20 min n = 7, S_C_ 80 min n = 5, L_E_ 20 min n = 5, L_E_ 80 min n = 4, with each individual contributing three regions (MB, OL, CB).

Following dissection, total RNA was extracted from the sample pools. Each pool was homogenized using a 2-mercaptoethanol lysing solution and a 21-gauge, 1 mL sterile syringe (BD; Franklin Lakes, NJ). RNA was extracted using the RNeasy Micro Kit (QIAGEN, Hilden, Germany), which included a DNase treatment step. Resulting RNA material was checked using gel electrophoresis with a 1% agarose gel to assure no genomic DNA contamination was present. Quality and relative quantity were assessed using a NanoDrop^®^ (Thermo Fisher Scientific, Waltham, MA), and resulting quantity measures were further verified using GloMax^®^ Luminometer (Promega, Madison, WI). Following extraction, aliquots of the samples were organized in a 96-well PCR plate and reverse transcribed to cDNA using the iScript™ Reverse Transcription Supermix kit and protocol (Bio-Rad Laboratories, Hercules, CA). The resulting 96-well plate with cDNA was used as source quantitative real-time polymerase chain reaction (qRT-PCR) analysis.

### Primer design

We used *Ribosomal Protein S5* (*rpS5*) as a reference gene (Evans, 2004; Evans, 2006; Evans and Wheeler, 2001), and three target genes: *CREB, CaMKII*, and *fen-1*. Reference gene primer sequences were obtained from previously published sources implementing rpS5 as a reference gene given its expression stability (Evans, 2004; Evans, 2006; Evans and Wheeler, 2001), which we also confirmed across tissue and time points (*data not shown*). The *fen-1* primers used in our study were previously developed and validated as part of the University of Puerto Rico at Rio Piedras 2010 Topicos Graduate Course (*data not shown*). For *CREB*, we developed primers specific to isoforms known to be specific in the brain (Eisenhardt et al., 2003).

*qRT PCR*. Optimized primer sets were used in conjunction with iTaq™ Universal SYBR^®^ Green Supermix (BioRad, Hercules, CA) and aliquots of our samples to conduct our qRT-PCR analysis. For each gene three 96-well PCR plates were run in a Stratagene™ MX3005P qPCR system. Resulting cycle thresholds (C_t_) were checked and samples that did not produce at least two consistent values across the three plates were discarded from the study. Replicates were discarded if the product T_m_ deviated by one degree from the expected amplicon and other resulting T_m_ values to avoid mis-priming or primer artifacts during amplification.

### Data analysis

We used −ΔΔCt to analyze resulting qRT-PCR expression data using the N_C_ group as the calibrator. Gene expression differences were examined gene-by-region between N_C_ and all other groups using a one-way ANOVA which combined treatment and timepoints into a single variable. This approach identified significant changes in expression related to experimental manipulation, with individual pairwise differences identified via a post-hoc Dunnett’s test using the N_C_ group as control. Significant changes in gene expression over time and between treatment groups were determined via a two-way ANOVA of C_C_, S_C_, and L_E_ groups and individual pairwise differences were identified via a Tukey’s post-hoc test. All statistical analyses were conducted using the R software (R Core Team, 2016).

## Results

Gene expression in the MB, where visual stimuli are processed and contextualized, showed dramatic changes both relative to N_C_ and across timepoints (Fig. 1). Each gene was significantly upregulated 80 min post-trial in the MB of L_E_ (*CaMKII*, t = 3.844, *p* = 0.003; *CREB*, t = 3.29, *p* = 0.01; *fen-1*, t = 2.02, *p* = 0.05). In addition, *CaMKII* and *fen-1* were significantly increased at 80 min relative to the 20 min timepoint (*CaMKII*, 20 vs. 80 min, t = −2.26, *p* = 0.0005; *fen-1*, 20 vs. 80 min, t = −2.11, *p* = 0.05). Our findings highlight that activation was observed earlier in the MB region and more pronounced across our target genes.

**Figure 1.**
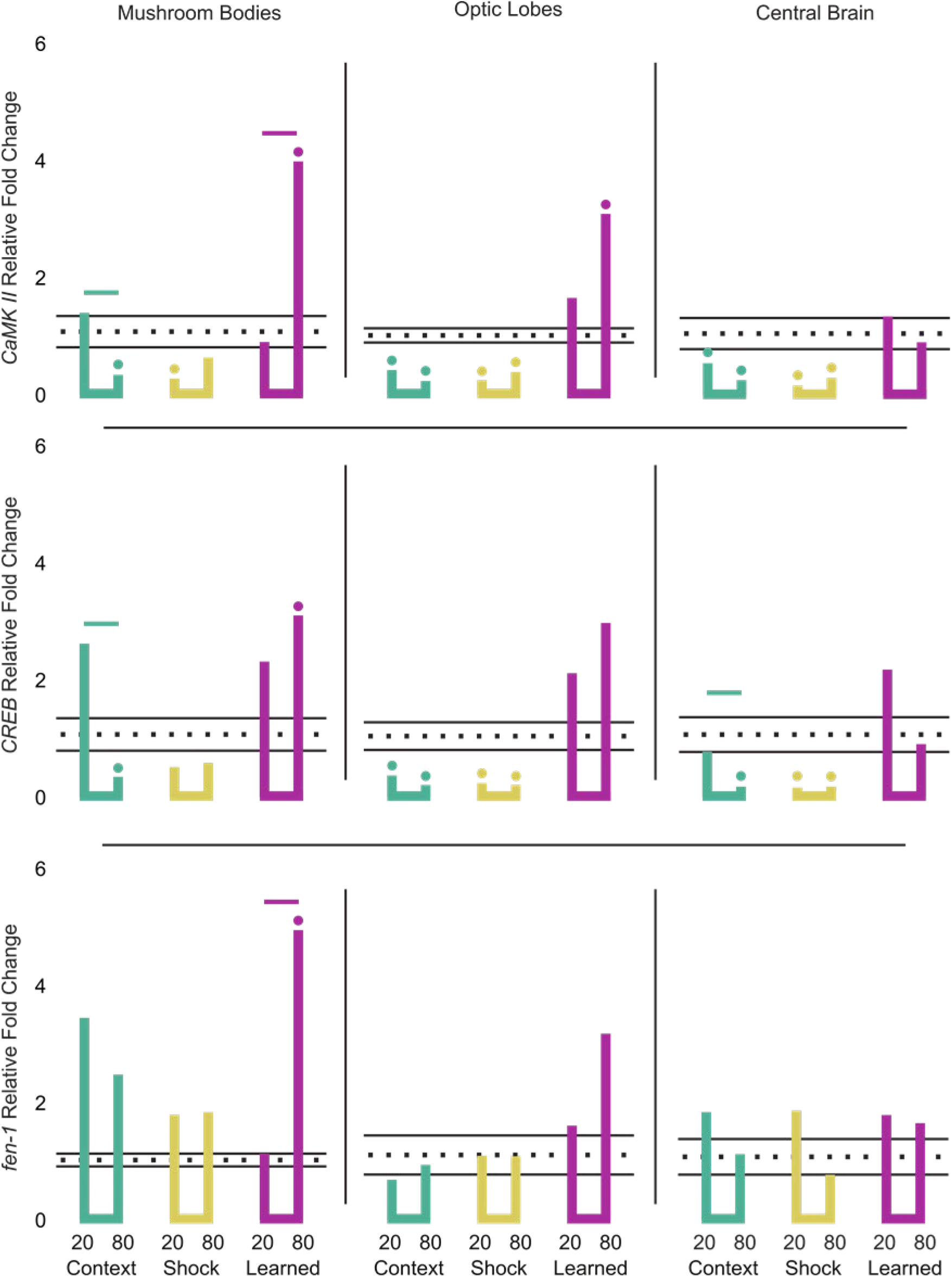
Expression profile of target genes following punishment learning. Each row in the figure corresponds to a target gene, each column to a broad brain region. In each panel section, the simplified bar graph shows lines extending towards the mean relative fold change in expression for each group and timepoint combination. The horizontal dotted line represents the mean relative fold change in expression of the N_C_ group with the solid lines delimiting the range within one standard error of the mean. Significant differences from N_C_ group expression are indicated by the circles atop the line. Significant expression level differences between timepoints within a group are identified by horizontal lines matching the group’s color.

Similarly, in the OL, where the initial process of sensory signal transduction occurs, we observed significant upregulation *CaMKII* in L_E_ at the 80 min time point (t = 3.899, *p* = 0.004; Fig. 1). Both *CREB* and *fen-1* were also elevated at 80 min, paralleling gene expression in the MB, but this relationship was not significant (Fig. 1). No other treatment group or timepoint showed a significant upregulation in this region, however both *CaMKII* and *CREB* were significantly downregulated in C_C_ and S_C_ relative to N_C_ at 20 (C_C_, t = −2.851, *p* = 0.04; S_C_, t = −6.154, *p* << 0.001) and 80 min post-trial (C_C_, t = −6.042, *p* << 0.001; S_C_, t = −4.006, *p* = 0.004). These results suggest learning-associated gene expression patterns in the OL that parallel those in the MB.

The CB region served as an experimental control as we do not anticipate that visual learning specifically activates the olfactory or gustatory system. The CB showed no significant upregulation though significant downregulation was observed in at the 80 min C_C_ group and both S_C_ groups relative to N_C_ group for *CaMKII* and *CREB* (*CaMKII*, C_C_ 80 min, t = −5.02, *p* < 0.001; *CaMKII*, S_C_ 20 min, t = −7.40, *p* < 0.001; *CaMKII*, S_C_ 80 min, t = −4.74, *p* < 0.001; CREB, C_C_ 80 min, t = −5.59, *p* < 0.001; CREB, S_C_ 20 min, t = −6.10, *p* < 0.001; CREB, S_C_ 80 min, t = −5.60, *p* < 0.001; Fig. 1). This signal suggests that though the regions aggregated in the CB are still responding to the exposure; it is a distinct response to those observed in either the OL or MB.

## Discussion

Though the MB has received considerable attention as the seat of learning and memory in the insect brain (Heisenberg, 1998; Heisenberg, 2003; Strausfeld, 2012), we show that a sensory neuropil, the OL, may also be a neuroanatomical substrate for learning, thus extending its known role beyond sensory transduction. This finding is evidence against our first hypothesis, which predicted that transcriptomic signal would be absent outside of the MB following a task that demanded learning. Rather, molecular signatures of learning and memory following aversive learning show similar patterns of gene expression in both the OL and MB, supporting parallel mechanisms of learning and memory in both tissues. This is supported by a recent study which investigated immediate early gene (IEG) expression in the honey bee brain and suggested that both sensory and higher-order brain regions express IEGs across a similar time course following an aggressive encounter (Traniello et al., 2019).

Interestingly, *CaMKII* upregulation following learning in both the MB and OL at 80 min, but not 20 min, post-learning implies a similar time course of activation across visual neuropil and higher-order processing centers in the honey bee brain. Furthermore, both *CREB* and *fen-1* showed increases in expression in the OL. Though nonsignificant, these responses were only seen at 80 min post-learning, further supporting collateral mechanisms which contrast with the predictions of Hypothesis 2. This gene expression time course further contrasts with spatiotemporal mapping of gene expression in larger vertebrate brains, where specific regions may be genetically activated relative to their place in a signal transduction pathway (Saul et al., 2019). We suggest that this difference may be related to spatial and metabolic constraints intrinsic to the arthropod brain that influence distinct processing strategies (Chittka and Niven, 2009; Niven and Farris, 2012).

In addition, our results show that *fen-1* expression increased in a region-specific manner. Initially, we considered that *fen-1* expression was not associated with learning but instead could be a response to shock-induced oxidative damage to DNA (Adachi et al., 1993; Lee et al., 2000). Our finding that *fen-1* was elevated in the MB and OL following aversive learning but not shock alone is evidence of a specific association with LTM. This suggests a conserved function of *fen-1* in aversive learning, previously described only in vertebrate systems (Saavedra-Rodríguez et al., 2009; Wang et al., 2003).

Our study relates neuroanatomical substrates to conserved molecular processes associated with visual memory formation. We show that distinct compartments of the honey bee brain are activated across a similar time course independent of their location in a neural circuit involved in learning. Further studies will be necessary to dissect peaks of upregulation for each gene in each region to determine if, for example, expression levels in the OL are comparable to those in the MB, but peak at distinct times following the learning assay. Here, we demonstrate involvement of a sensory region not typically associated with learning, thus implying that the memory of environmental experience is distributed across distinct anatomical regions of the honey bee brain.

**Table 1.**
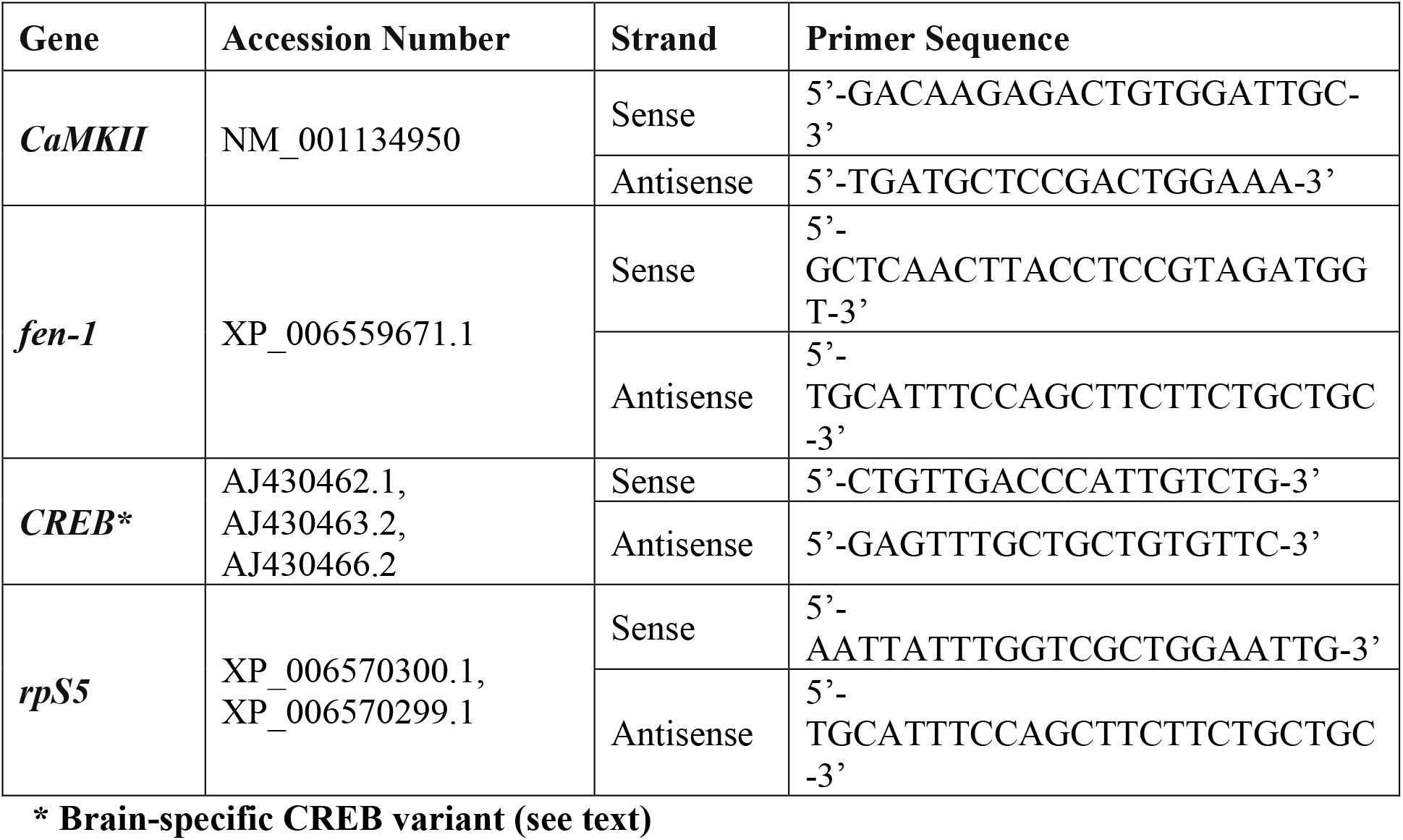
Primer sets used for each of the target candidate genes. This table reports primer sequences for the panel of genes used. Specifically provided are the genes’ name, GenBank accession number, primer strand read direction, and actual sequence.

## Acknowledgements

This work was supported by a National Institute for General Medical Sciences RISE Graduate Fellowship (R25GM061151-11) to A.A., a National Science Foundation Postdoctoral Fellowship (Program 15-501) to A.A, a National Science Foundation OISE award (#1545803) to T.G., and a Catalyzer Research Grant from the Puerto Rico Science Technology Research Trust to T.G.

## Tables and Figures

**Supplementary Figure 1.**
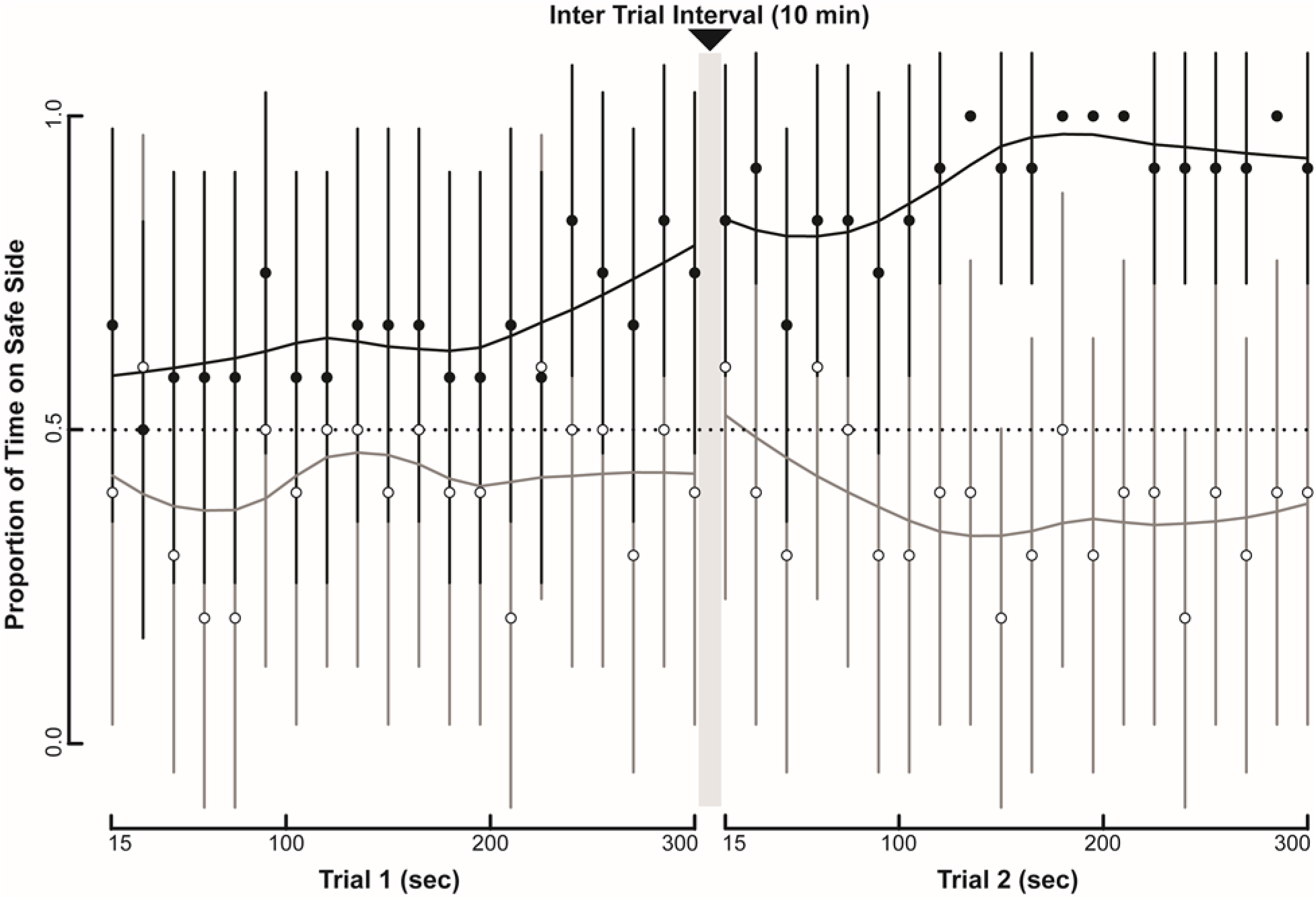
Summary of behavioral response used for sample selection. The plot represents learning response of test bees. Dark circles identify the group mean for honey bees that met sample selection criteria and were included in the L_E_ (n = 18), light circles identify cohorts that did not meet selection criteria and were not considered (n = 8, *see Methods*). Vertical dark and light bars represent the 95% confidence interval for each corresponding-colored group. A Loess smoothing curve is also provided to visualize the learning trend for each group and each Trial.

**Supplementary Figure 2.**
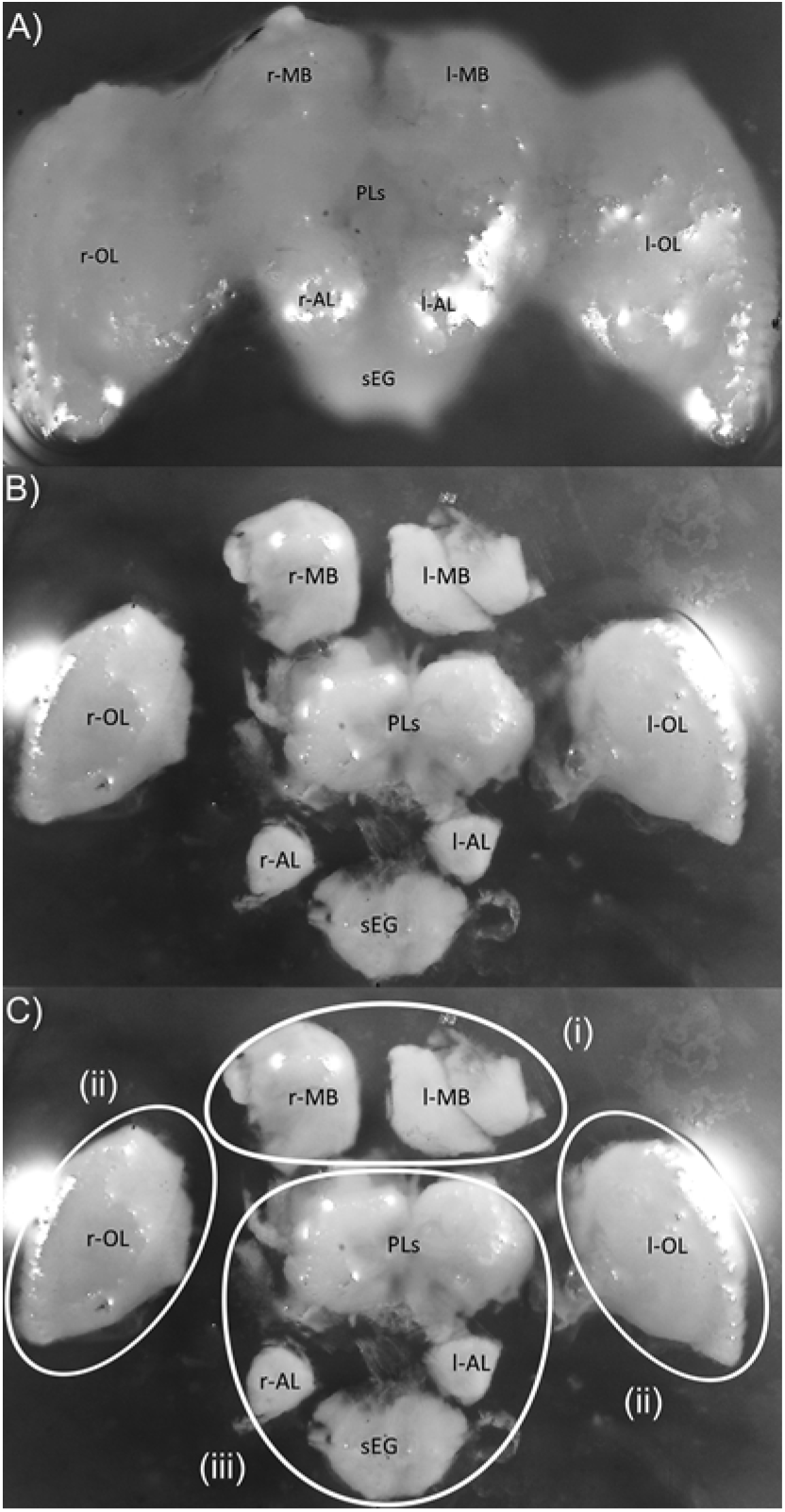
Dissected and sub-sectioned honey bee brain defining broad brain regions analyzed via qRT-PCR Methods. The figure demonstrates a dissected (A) honey bee worker brain with annotated gross anatomical regions that include right and left mushroom bodies (r-, l-MB), proto-cerebrum (PC), right and left optic lobes (r-, l-OL), right and left antennal lobes (r-, l-AL), and the subesophageal ganglion (sEG). Further depicted is the same brain, sub-sectioned into these gross anatomical regions (B), and later those same gross anatomical regions highlighted (C) to denote the three broad brain regions used in the study, namely: (i) mushroom bodies (MB), (ii) optic lobes (OL), and (iii) central brain (CB).

## References

Adachi, S., Kawamura, K. and Takemoto, K. (1993). Oxidative damage of nuclear DNA in liver of rats exposed to psychological stress. Cancer Res. 53, 4153–4155.

Agarwal, M., Giannoni Guzmán, M., Morales-Matos, C., Del Valle Díaz, R. A., Abramson, C. and Giray, T. (2011). Dopamine and octopamine influence avoidance learning of honey bees in a place preference assay. PLoS One 6, e25371.

Avalos, A., Pérez, E., Vallejo, L., Pérez, M. E., Abramson, C. I. and Giray, T. (2017). Social signals and aversive learning in honey bee drones and workers. Biol. Open 6, 41–49.

Bito, H., Deisseroth, K. and Tsien, R. W. (1996). CREB phosphorylation and dephosphorylation: A Ca2+- and stimulus duration-dependent switch for hippocampal gene expression. Cell 87, 1203–1214.

Bitterman, M. E., Menzel, R., Fietz, A. and Schäfer, S. (1983). Classical conditioning of proboscis extension in honeybees (Apis mellifera). J. Comp. Psychol. 97, 107–119.

Brandt, R., Rohlfing, T., Rybak, J., Krofczik, S., Maye, A., Westerhoff, M., Hege, H.-C. and Menzel, R. (2005). Three-dimensional average-shape atlas of the honeybee brain and its applications. J. Comp. Neurol. 492, 1–19.

Cartwright, B. A. and Collett, T. S. (1983). Landmark learning in bees. J. Comp. Physiol. A 151, 521–543.

Chittka, L. and Niven, J. (2009). Are Bigger Brains Better? Curr. Biol. 19, R995–R1008.

Chittka, L. and Raine, N. E. (2006). Recognition of flowers by pollinators. Curr. Opin. Plant Biol. 9, 428–435.

Eisenhardt, D., Friedrich, a, Stollhoff, N., Müller, U., Kress, H. and Menzel, R. (2003). The AmCREB gene is an ortholog of the mammalian CREB/CREM family of transcription factors and encodes several splice variants in the honeybee brain. Insect Mol. Biol. 12, 373–82.

Evans, J. (2004). Transcriptional immune responses by honey bee larvae during invasion by the bacterial pathogen, Paenibacillus larvae. J. Invertebr. Pathol. 85, 105–111.

Evans, J. (2006). Beepath: An ordered quantitative-PCR array for exploring honey bee immunity and disease. J. Invertebr. Pathol. 93, 135–9.

Evans, J. and Wheeler, D. E. (2001). Expression profiles during honeybee caste determination. Genome Biol. 2, RESEARCH0001.

Giurfa, M., Núñez, J., Chittka, L. and Menzel, R. (1995). Colour preferences of flower-naive honeybees. J. Comp. Physiol. A 177, 247–259.

Heisenberg, M. (1998). What do the mushroom bodies do for the insect brain? An introduction. Learn. Mem. 5, 1–10.

Heisenberg, M. (2003). Mushroom body memoir: From maps to models. Nat. Rev. Neurosci. 4, 266–275.

Kamikouchi, A., Takeuchi, H., Sawata, M., Natori, S. and Kubo, T. (2000). Concentrated expression of Ca2+/ Calmodulin-Dependent Protein Kinase II and Protein Kinase C in the mushroom bodies of the brain of the honeybee Apis mellifera L. J. Comp. Neurol. 417, 501–10.

Kandel, E. (2001). The molecular biology of memory storage: A dialogue between genes and synapses. Science 294, 1030–8.

Kandel, E. (2012). The molecular biology of memory: cAMP, PKA, CRE, CREB-1, CREB-2, and CPEB. Mol. Brain 5, 14.

Lakhina, V., Arey, R. N., Kaletsky, R., Kauffman, A., Stein, G., Keyes, W., Xu, D. and Murphy, C. T. (2015). Genome-wide functional analysis of CREB/long-term memory-dependent transcription reveals distinct basal and memory gene expression programs. Neuron 85, 330–345.

Lee, R. C., Zhang, D. and Hannig, J. (2000). Biophysical injury mechanisms in electrical shock trauma. Annu. Rev. Biomed. Eng. 2, 477–509.

Lehrer, M., Horridge, G. A., Zhang, S. W. and Gadagkar, R. (1995). Shape vision in bees: Innate preference for flower-like patterns. Philos. Trans. R. Soc. B Biol. Sci. 347, 123–137.

Matsumoto, Y. and Mizunami, M. (2002). Temporal determinants of long-term retention of olfactory memory in the cricket Gryllus bimaculatus. J. Exp. Biol. 205, 1429–37.

Matsumoto, Y., Hatano, A., Unoki, S. and Mizunami, M. (2009). Stimulation of the cAMP system by the nitric oxide-cGMP system underlying the formation of long-term memory in an insect. Neurosci. Lett. 467, 81–5.

Mattu, V., Raj, H. and Thakur, M. (2012). Foraging behavior of honeybees on apple crop and its variation with altitude in Shimla Hills of Western Himalaya, India. Int. J. Sci. Nat. 3, 296–301.

Menzel, R. (1999). Memory dynamics in the honeybee. J. Comp. Physiol. A Sensory, Neural, Behav. Physiol. 185, 323–340.

Menzel, R. and Giurfa, M. (2001). Cognitive architecture of a mini-brain: The honeybee. Trends Cogn. Sci. 5, 62–71.

Menzel, R., Greggers, U., Smith, A., Berger, S., Brandt, R., Brunke, S., Bundrock, G., Hulse, S., Plumpe, T., Schaupp, F., et al. (2005). Honey bees navigate according to a map-like spatial memory. Proc. Natl. Acad. Sci. 102, 3040–3045.

Niven, J. E. and Farris, S. M. (2012). Miniaturization of nervous systems and neurons. Curr. Biol. 22, R323–R329.

Nouvian, M. and Galizia, C. G. (2019). Aversive training of honey bees in an automated y-maze. Front. Physiol. 10, 1–17.

Pasch, E., Muenz, T. S. and Rössler, W. (2011). CaMKII is differentially localized in synaptic regions of Kenyon cells within the mushroom bodies of the honeybee brain. J. Comp. Neurol. 519, 3700–12.

Perisse, E., Raymond-Delpech, V., Néant, I., Matsumoto, Y., Leclerc, C., Moreau, M. and Sandoz, J.-C. (2009). Early calcium increase triggers the formation of olfactory long-term memory in honeybees. BMC Biol. 7, 30.

R Core Team (2016). R: A language and environment for statistical computing.

Rybak, J., Kuß, A., Lamecker, H., Zachow, S., Hege, H.-C., Lienhard, M., Singer, J., Neubert, K. and Menzel, R. (2010). The digital bee brain: Integrating and managing neurons in a common 3D reference system. Front. Syst. Neurosci. 4,.

Saavedra-Rodríguez, L., Vázquez, A., Ortiz-Zuazaga, H. G., Chorna, N. E., González, F. A., Andrés, L., Rodríguez, K., Ramírez, F., Rodríguez, A. and Peña de Ortiz, S. (2009). Identification of flap structure-specific endonuclease 1 as a factor involved in long-term memory formation of aversive learning. J. Neurosci. 29, 5726–5737.

Saul, M. C., Blatti, C., Yang, W., Bukhari, S. A., Shpigler, H. Y., Troy, J. M., Seward, C. H., Sloofman, L., Chandrasekaran, S., Bell, A. M., et al. (2019). Cross-species systems analysis of evolutionary toolkits of neurogenomic response to social challenge. Genes, Brain Behav. 18, e12502.

Seeley, T. D. and Visscher, P. K. (2003). Choosing a home: How the scouts in a honey bee swarm perceive the completion of their group decision making. Behav. Ecol. Sociobiol. 54, 511–520.

Sen Sarma, M., Rodriguez-Zas, S. L., Gernat, T., Nguyen, T., Newman, T. and Robinson, G. E. (2010). Distance-responsive genes found in dancing honey bees. Genes, Brain Behav. 9, 825–830.

Strausfeld, N. J. (2002). Organization of the honey bee mushroom body: Representation of the calyx within the vertical and gamma lobes. J. Comp. Neurol. 450, 4–33.

Strausfeld, N. J. (2012). Arthropod brains: Evolution, functional elegance, and histroical significance. Cambridge: Belknap Press of Harvard University Press.

Traniello, I. M., Chen, Z., Bagchi, V. A. and Robinson, G. E. (2019). Valence of social information is encoded in different subpopulations of mushroom body Kenyon cells in the honeybee brain. Proc. R. Soc. B Biol. Sci. 286, 20190901.

Visscher, P. K. (2007). Group decision making in nest-site selection among social insects. Annu. Rev. Entomol. 52, 255–275.

Wang, J., Ren, K., Pérez, J., Silva, A. J. and Peña de Ortiz, S. (2003). The antimetabolite ara- CTP blocks long-term memory of conditioned taste aversion. Learn. Mem. 10, 503–509.

Witthöft, W. (1967). Absolute anzahl und verteilung der zellen im him der honigbiene. Zeitschrift für Morphol. der Tiere 61, 160–184.

